# Microscale tracking of coral disease reveals timeline of infection and heterogeneity of polyp fate

**DOI:** 10.1101/302778

**Authors:** Assaf R. Gavish, Orr H. Shapiro, Esti Kramarsky-Winter, Assaf Vardi

## Abstract

Coral disease is often studied at scales ranging from single colonies to the entire reef. This is particularly true for studies following disease progression through time. To gain a mechanistic understanding of key steps underlying infection dynamics, it is necessary to study disease progression, and host-pathogen interactions, at relevant microbial scales. Here we provide a dynamic view of the interaction between the model coral pathogen *Vibrio coralliilyticus* and its coral host *Pocillopora damicornis* at unprecedented spatial and temporal scales. This view is achieved using a novel microfluidics-based system specifically designed to allow microscopic study of coral infection *in*-*vivo* under controlled environmental conditions. Analysis of exudates continuously collected at the system’s outflow, allows a detailed biochemical and microbial analyses coupled to the microscopic observations of the disease progression. The resulting multilayered dataset provides the most detailed description of a coral infection to-date, revealing distinct pathogenic processes as well as the defensive behavior of the coral host. We provide evidence that infection in this system occurs following ingestion of the pathogen, and may then progress through the gastrovascular system. We further show infection may spread when pathogens colonize lesions in the host tissue. Copious spewing of pathogen-laden mucus from the polyp mouths results in effective expulsion of the pathogen from the gastrovascular system, possibly serving as a first line of defense. A secondary defense mechanism entails the severing of calicoblastic connective tissues resulting in the controlled isolation of diseased polyps, or the survival of individual polyps within infected colonies. Further investigations of coral-pathogen interactions at these scales will help to elucidate the complex interactions underlying coral disease, as we as the versatile adaptive response of the coral ecosystems to fluctuating environments.

## Introduction

Coral reefs are currently undergoing an unprecedented decline driven by local and global changes to their environment^1^. Reef building corals, commonly described as holobionts, form a complex relationship with photosynthesizing dinoflagellates (*Symbiodinium spp*.) and a consortium of microbial partners^2^. Shifts in environmental conditions may lead to the breakdown of these symbiotic relations, often with catastrophic consequences for the coral colony. Such processes, collectively termed coral disease^3, 4^, may be manifested as a loss of the algal symbionts (coral bleaching)^5^, or as damage to the coral colony due to various forms of necrotic loss of coral tissue^2^. On large scales, these processes may result in loss of coral cover, ultimately leading to the degradation of the reef structure and the loss of associated ecological and societal services^4, 6, 7^.

Many coral diseases are linked to specific pathogens whose abundance and virulence increase in response to environmental changes. Such changes may include nutrient loading, pollution, and temperature shifts^8-11^. One of the best characterized coral diseases is the infection of the Indo-Pacific coral *Pocillopora damicornis* by the bacterial pathogen *Vibrio coralliilyticus*^9, 12^, ^13^. The virulence of *V. coralliilyticus* is known to be positively correlated with increased temperatures^9^, ^14-16^. Increased ambient temperatures are further linked to accelerated vibrio growth rates^9^, enhanced chemotaxis and chemokinesis^17^, and secretion of matrix metalloproteases (MMPs)^18^. Nevertheless, a mechanistic understanding linking these traits to coral infection and disease progress is still lacking.

Many coral disease studies focus on monitoring coral colonies for the appearance of macroscopic signs of disease. These may include various forms of tissue discoloration, loss of the algal symbionts, or loss of tissue integrity^19, 20^. This tendency for macroscale studies is derived to a large extent from the complexity of the coral holobiont^21, 22^, and the difficulty in establishing a tractable model system facilitating more detailed observations^22, 23^. Currently, the main available tool enabling to link a potential pathogen to the site of tissue damage and to the host response is histopathology^23^. However, as such disease manifestations only appear at advanced stages of the infection process, their use as disease indicators fails to capture the early stages of pathogen colonization and disease initiation^23^. We are thus lacking a mechanistic understanding of key steps in the infection process, including e.g. site of initial colonization, possible functions of specific disease markers such as MMP’s, or where bacterial chemotaxis may come into play. Furthermore, there are still major knowledge gaps in our understanding of coral response at the onset of pathogenic infection.

Here we present a new microfluidic system, the Microfluidic Coral Infection (MCI) platform, developed specifically to tackle question related to the interaction between a bacterial pathogen and a coral colony at high spatio-temporal resolutions. This platform has several features distinguishing it from the previously published coral-on-a-chip (CoC) system^24^. The larger chamber volume and higher flow rates of the MCI, as compared to the CoC, facilitate the incubation of small coral fragments, preserving the colonial morphology of the coral colony. Moreover, the MCI design allows the continuous collection of exudates of the system for downstream analysis. The MCI further allows the incubation and tracking of up to 6 individual coral fragments in separate chambers, facilitating flexible experimental design. Using the MCI system we track the microscopic encounter between the bacterial pathogen *V. coralliilyticus* and its coral host *P. damicornis.* Coupling the resulting time-lapse microscopic imaging with biochemical and microbial analyses of the system’s outflow we identify early stages of the infection process that were not previously described. These results bring us a step closer towards a mechanistic understanding of microbial disease processes in reef building corals.

## Results

### Live imaging of coral infection

The progression of bacterial infection of small coral fragments was enabled using the MCI platform (Figure 1; Supplementary figure 1). To demonstrate the robustness of this system, healthy *P. damicornis* fragments were incubated under controlled environmental conditions (temperature, light, flow, and water quality). Tissue integrity was continuously monitored by microscopic imaging. Natural coral Green Fluorescence Protein (GFP) served as a biomarker for coral health, while chlorophyll autofluorescence served to track localization and wellbeing of its zooxanthellae symbionts. No changes in coral morphology or behavior, and no extensive loss of algal symbionts (bleaching) were observed following 48 h of incubation under constant flow of filtered artificial seawater (FASW).

**Figure 1:**
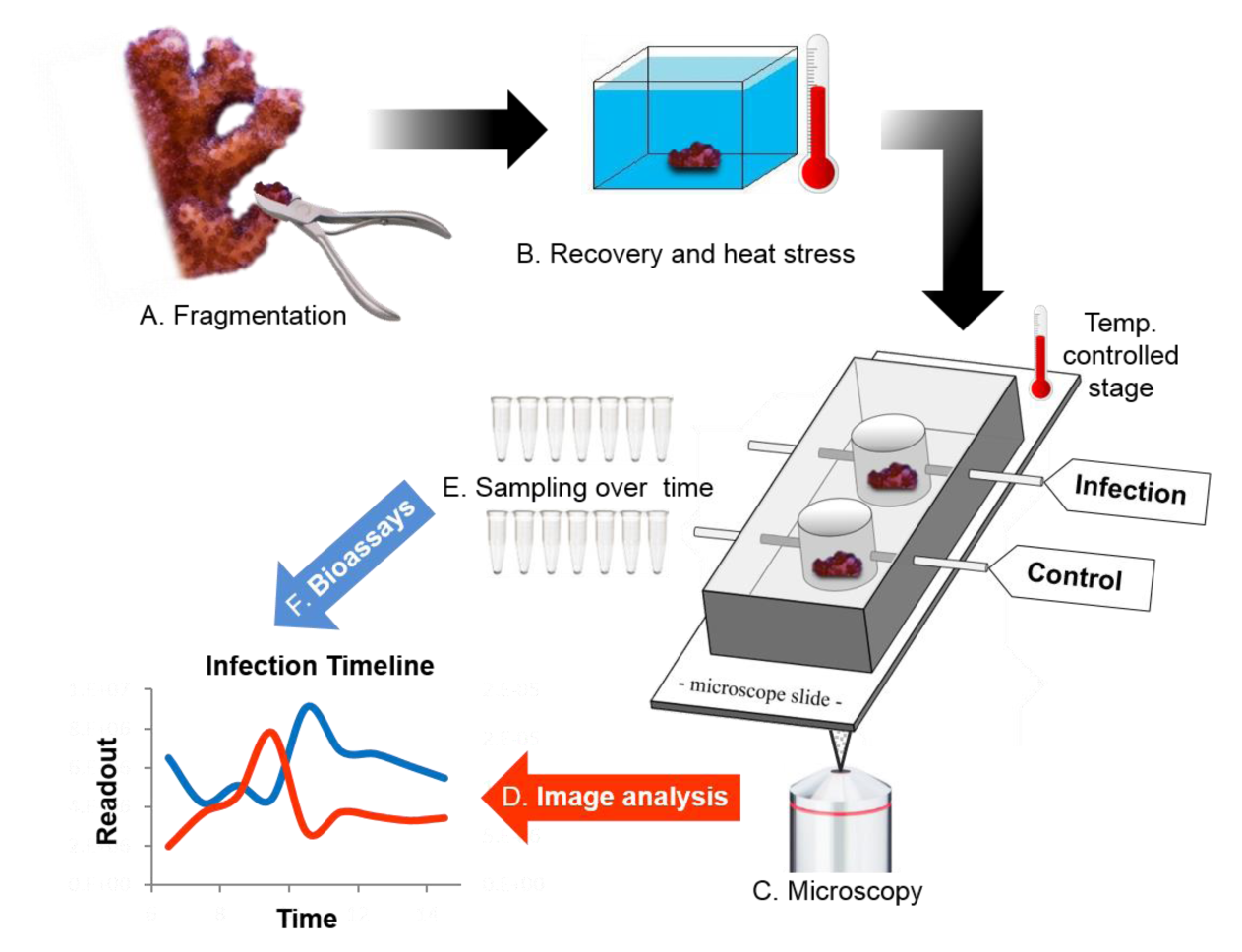
Experimental work-flow for Microfluidic Coral Infection (MCI). A, B. Small coral fragments (~3-5 mm), clipped from the branch tips of a *P. damicornis* colony, are kept in the main tank for recovery. Fragments are incubated for 3 days in a small (3.5 L) temperature-controlled tank filled with filtered aquarium water. Heat stress is induced by setting tank temperature to 30°C. **C**. Fragments are transferred to the MCI device placed on a temperature-controlled microscope stage. Infection is initiated by introducing DsRed-labelled *V. coralliilyticus* cells at desired duration and concentration through the inflow. Infection progress is tracked using epifluorescence and light microscopy at set intervals. **D**. Image analysis is used to quantify signal intensity and localization in all channels throughout the infection period. **E**. An automated fraction collector is used to sample flow through at set intervals throughout the experiment. Collected fractions are immediately cooled to below 2°C, with or without addition of fixative, for subsequent analysis. **F**. Collected fractions are analyzed by various bioassays, enabling correlation of microscopic observations and downstream analysis.

During infection experiments, challenged coral fragments were inoculated with either the bacterial pathogen *V. coralliilyticus* or the non-pathogenic *V. fischeri*, both labeled by DsRed^24^, ^25^ to facilitate imaging. Non-challenged fragments received FASW throughout the experiment. Tissue integrity, coral behavior, and zooxanthellae fluorescence and localization, as well as localization of labelled bacteria in challenged fragments, were microscopically monitored throughout each experiment (Figure 2).

**Figure 2:**
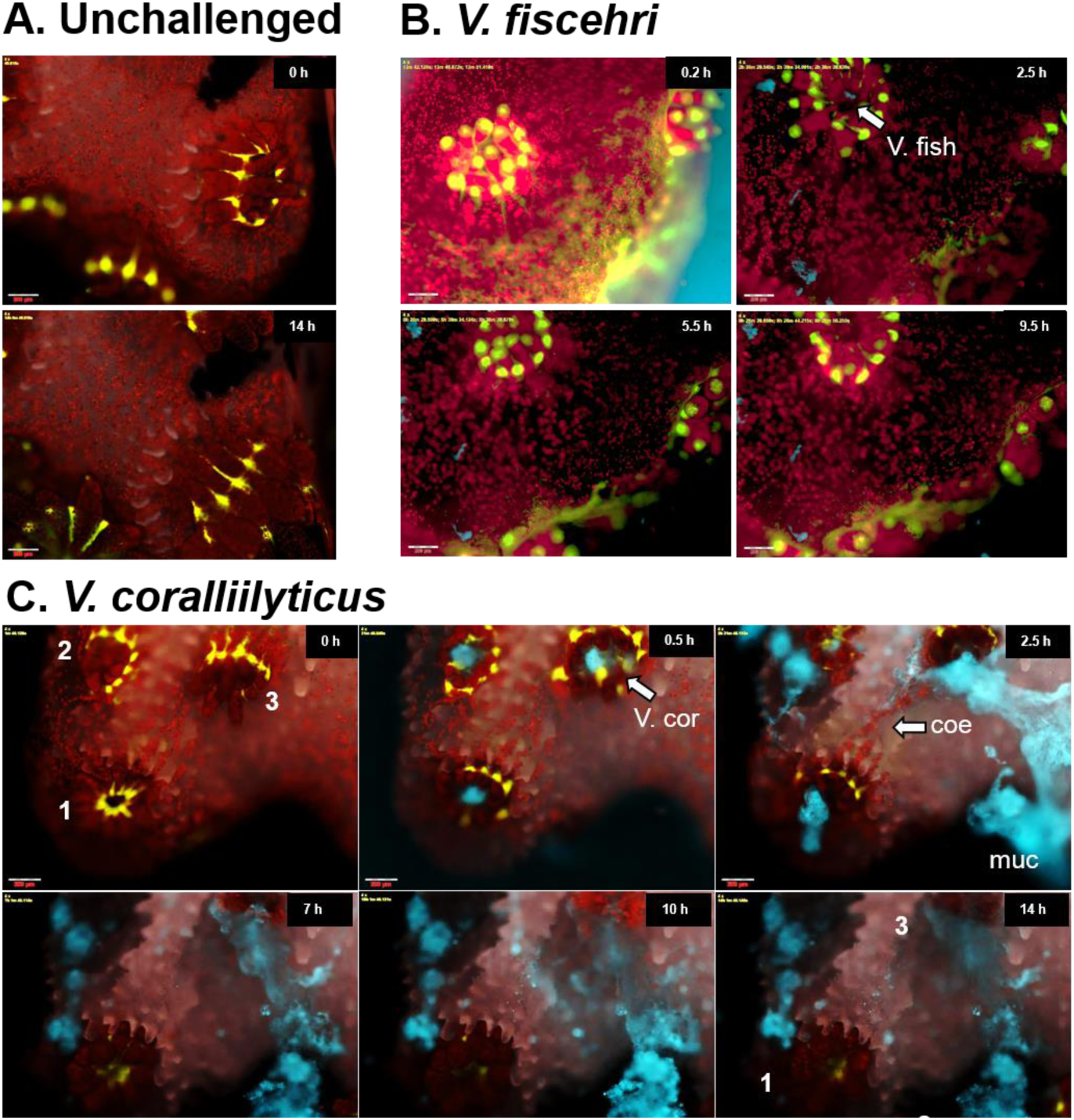
Timeline of a coral infection. A. Microscopic view of an unchallenged control *P. damicornis* fragment showing coral GFP (Green) and algal chlorophyll (Red). All nonchallenged fragments appeared healthy at the end of the experiment (here 14 hours), with tentacles extended and no apparent bleaching or tissue loss. **B.** In *P. damicornis* fragments challenged by *V. fischeri*, slight accumulation of DsRed-labelled bacteria (cyan; V. fish) was observed in the coral pharynx. No disease-like symptoms were observed. **C.** Fragments challenged by *V. coralliilyticus* regularly displayed behavioral and morphological changes: Time **‘0 h’**-Immediately prior to inoculation. Image shows three polyps (1-3) with partially extended tentacles. **0.5 h** – DsRed-labelled *V. coralliilyticus* (Cyan) accumulate at the polyp pharynx, but not on other exposed areas of the colony. **2.5 h** – Polyps secrete large amounts of mucus (muc), clearly visible due to large numbers of *V. coralliilyticus* cells adhering to it. Tearing of coenosarc tissue (coe) is observed. **7 h** –Coenosarc is degraded and polyps separated. Polyp **2** underwent polyp bail-out and is lost from the field of view. Polyp **3** is dead, with tissue degraded and GFP signal lost, although some chlorophyll autofluorescence is still observed. **10 h** - *V. coralliilyticus* accumulates on the exposed skeleton. **14 h** –At the end of the experiment polyp **1** remains viable, despite complete loss of surrounding tissue. A bailedout polyp (possibly polyp **2**) is visible at the bottom of the image. Vacant calyx of polyp **3** is marked. Scale for all images are 200 μm.

No morphological or behavioral changes were observed in non-challenged control fragments from all experiments (Fig. 2A; supplementary video 1). In fragments challenged by *V. fischeri* (10^8^ cells/ml), accumulation of DsRed-labelled bacteria was observed in the polyp pharynx over the 1^st^ hour of inoculation. This was followed by moderate spewing of bacterial-laden mucus from all polyps (Fig 2B; supplementary video 2). No other morphological or behavioral changes were observed.

A markedly different response was observed in fragments challenged by *V. coralliilyticus* (10^8^ cells/ml). A total of 39 fragments, derived from 6 coral colonies, were challenged over the course of 18 separate experiments (Supplementary Table 1). Within 15 minutes of inoculation, polyp contraction was observed in all *V. coralliilyticus*-*challenged* fragments. Pathogens accumulated in the coral pharynx over the next hour, with little or no accumulation observed on other parts of the coral surface (Figure 2C). Over the following 2-3 hours, polyps released copious amounts of viscous, bacterial-laden mucus, concomitant with substantial stretching of the coenosarc tissue connecting neighboring polyps (Figure 2C, Supplementary Video 3, 4).

Following this stage, experiment results followed one of two outcomes (Figure 3). In 7 of the challenged experiments, consisting of 14 coral fragments, a complete or near-complete recovery was observed (Supplementary Table 1). In these fragments tissue confluence was retained, and within a few hours of inoculation polyps expanded, with no labeled *V. coralliilyticus* cells observed in the pharynx (Supplementary video 3).

**Figure 3:**
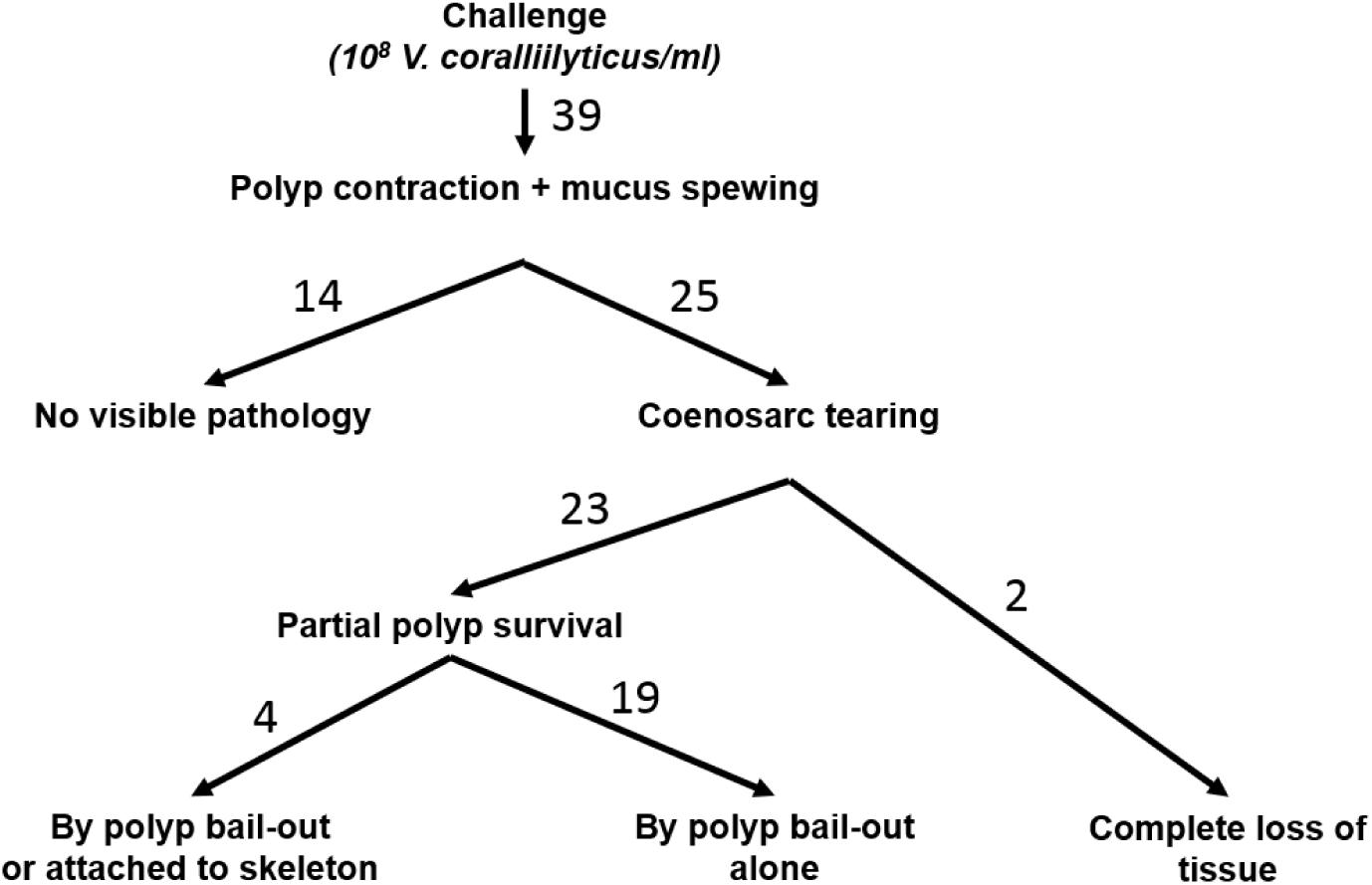
A roadmap of outcomes of for 39 *P. damicornis* fragments challenged by *V. coralliilyticus.* Polyp retraction and subsequent mucus spewing was observed in all fragments. In symptomatic fragments, this was followed by separation of neighboring polyps through coenosarc tearing. Individual polyps underwent one of three different fates (survival, bail-out or death).

Contrastingly, in the remaining 11 experiments, consisting of 25 challenged fragments, a clear pathology was observed (Figure 2C; Supplementary video 4). In these fragments, mucus spewing was followed by tearing of the coenosarc, leading to the separation of neighboring polyps and consequently loss of colony integrity (Fig. 2C). The majority of polyps in these experiments then underwent necrosis, manifested as visible loss of tissue integrity, accompanied by a gradual decay in GFP fluorescence (Figure 2C [Polyp 3]; Figure 4B; Supplementary figure 2). In 23 of these 25 fragments partial survival was observed in the form of polyp bail-out^24, 26^. In 4 of these 23 fragments, individual polyps survived and remained attached to the skeleton at the end of the experiment (Figure 2C; Figure 3; Supplementary Video 4; Supplementary Table 1).

**Figure 4:**
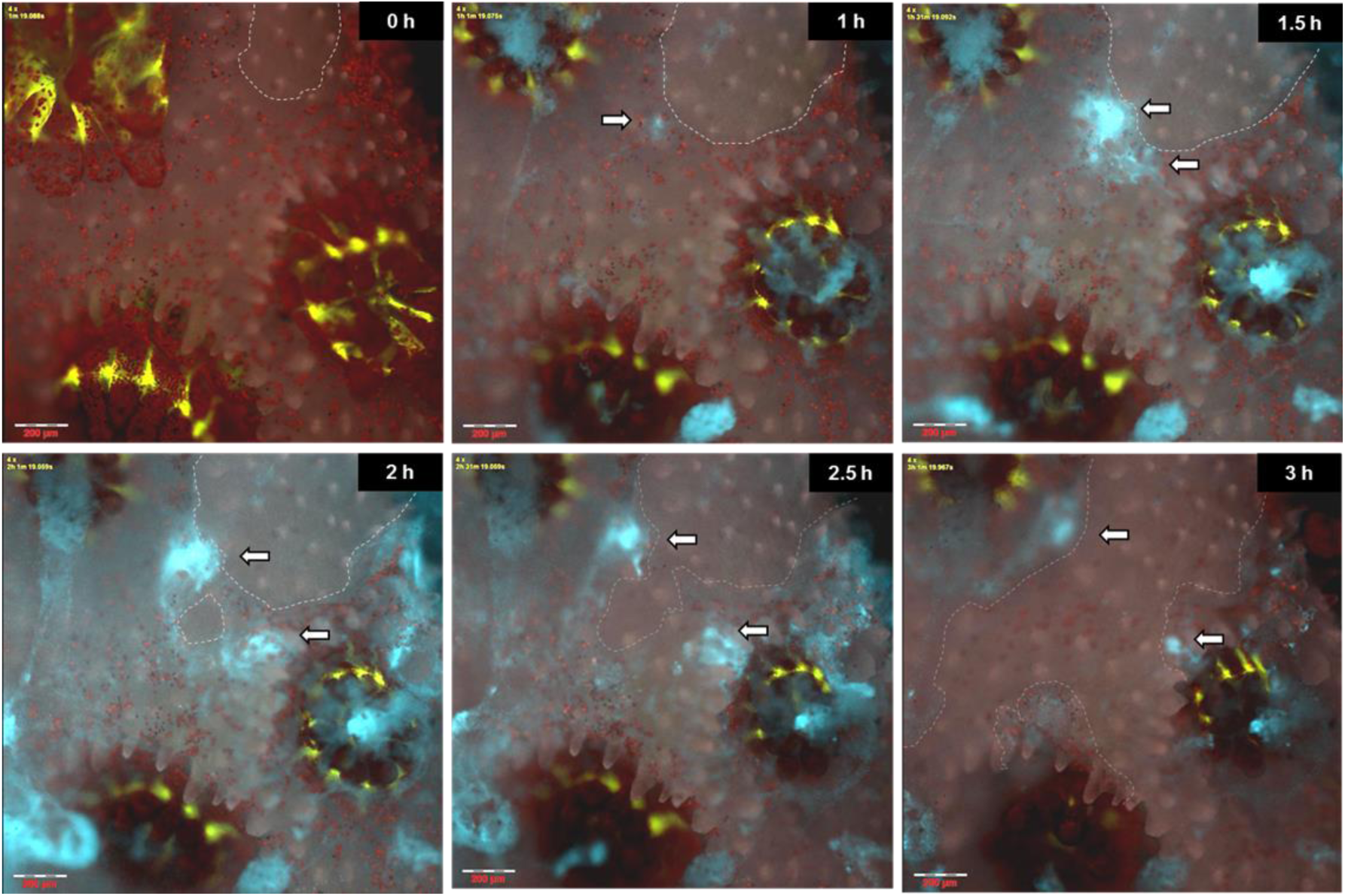
Tissue lesions allow rapid colonization and infection. Time **‘0’ h** – Infected fragmented immediately prior to inoculation. Coral GFP (Green) and algal chlorophyll (Red) are shown on a greyscale background. Dashed line marks the borders of a small lesion approximately 300 μm in diameter. **1 h** – *V. coralliilyticus* cells (Cyan) accumulate at the lesion edge (arrow). **1.5 h – 3 h** – further colonization of the torn tissue is followed by rapid lesion expansion and death of neighboring polyps. The complete sequence from this experiment is provided in supplementary video 4. Scale bars are 200 μm.

### Lesion infection

A slightly modified infection sequence was observed in fragments with minor lesions in the coenosarc tissues. *V. coralliilyticus* cells regularly accumulated at the lesion edge (Figure 4; Supplementary Video 5) within the first hour of inoculation. Further bacterial accumulation or proliferation was observed over the next hours, accompanied by tissue necrosis manifested as rapid tearing or degradation of the ceonosarc tissue and death of neighboring polyps.

### Quantitative image analysis

Quantification of fluorescence intensity derived from the different components of the holobiont (GFP for coral tissue, chlorophyll for algal symbionts and DsRed for *V. coralliilyticus*) provided further information on the infection progression in challenged fragments (Figure 5A, B). A gradual but constant decrease in coral GFP intensity, beginning approximately 2 h post inoculation, was consistently observed in dying fragments, but not in non-challenged controls or in challenged, asymptomatic fragments (Figure 5A, B). The most prominent feature in the resulting DsRed intensity profile was a large peak spanning the first 2 hours of each experiment, reflecting the inflow of labeled *V. coralliilyticus* during inoculation (Figure 4B). A smaller peak in the DsRed channel regularly appeared between 6 and 10 hours following inoculation. No distinct patterns were observed in chlorophyll autofluorescence within the timeframe of the infections described here.

**Figure 5:**
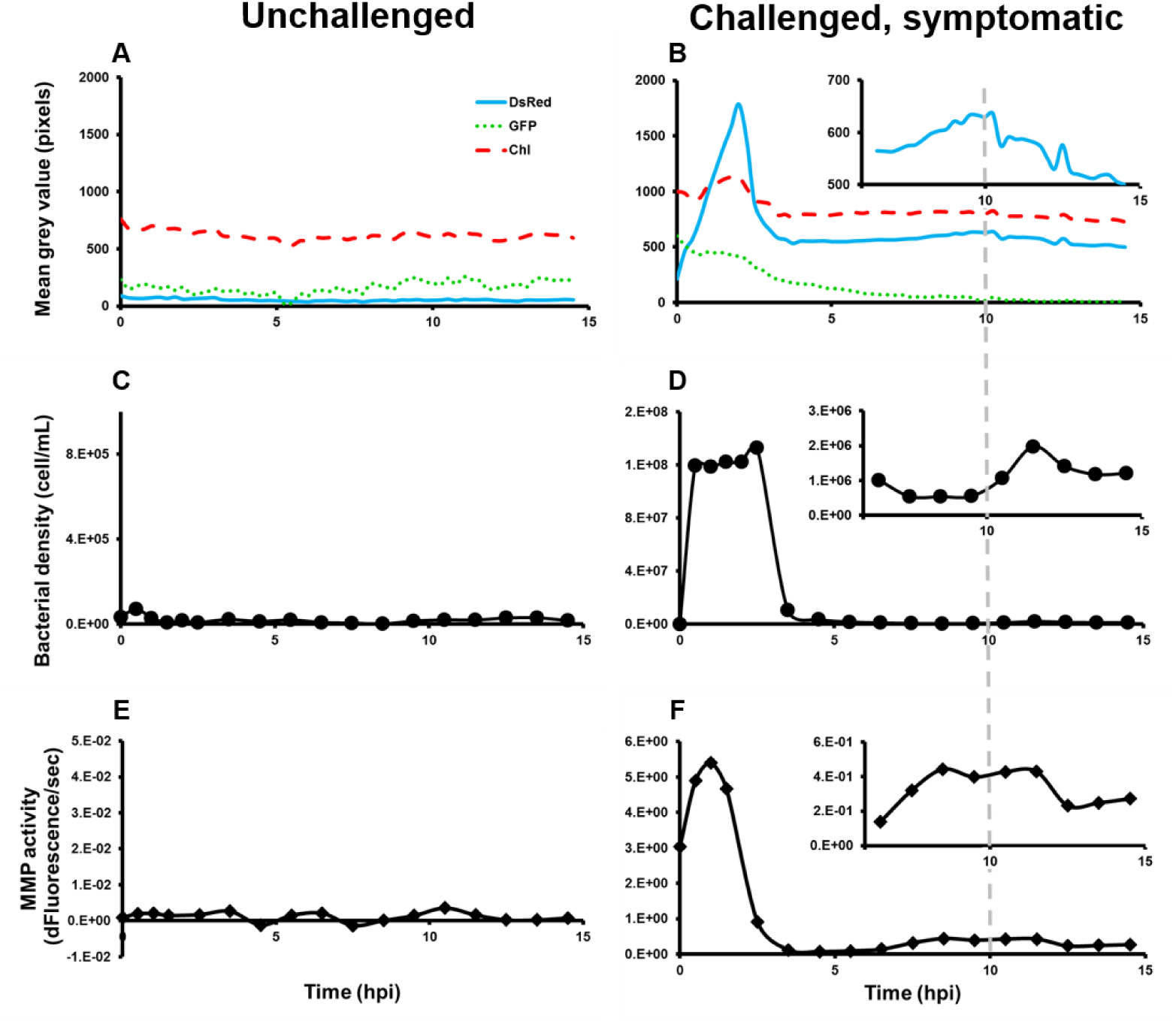
Quantitative analysis of microscope images and system exudates provide further insights into the timeline of coral infection. **A**, **B** – Quantification of fluorescence signals from GFP (green), Chlorophyll (red) and DsRed labeled *V. coralliilyticus* (cyan) in unchallenged (**A**) and challenged by *V. coralliilyticus* (**B**) fragments. High levels of DsRed signal over the first 2 hours of experiment in **B** correspond to pathogen inoculation and settlement of the coral pharynx. A gradual decrease in GFP signal starting approximately 2 hours post-infection indicates death and disintegration of the coral host. The increase in DsRed signal between 6 and 10 h (**B**, inset) likely indicates pathogen proliferation at the expense of the dying coral. **C, D**. Quantification of total bacterial density in the outflow of a control (**C**) and infected (**D**) chambers. High bacterial load in **D** over the first 2 hours indicate relatively low attachment of inoculated *V. coralliilyticus* to the coral host. A slight increase in bacterial density starting at 10 h (**D**, inset), termed “late burst”, is likely driven by pathogens, and possibly other bacteria, released from the dying tissue. **E, F**. Quantification of matrix metalloproteinases (MMPs) activity in the outflow of a control (**E**) and infected (**F**) chambers. Initial high activity corresponds to MMPs activity in the inoculum. Increased activity starting at 7 hours (**F**, inset) may indicate increased MMPs production by *V. coralliilyticus* as it breaks down the host tissue.

### Downstream microbial and biochemical analysis of MCI exudates

Additional insight into the infection process was gained by time-resolved measurement of microbial abundance and MMP activity in the MCI exudates (Figure 5C-F; Supplementary figures 3, 4). The highest values for both MMP activity and cell abundance were measured during the initial two-hour inoculation period. In all challenged fragments, bacterial abundance in the exudates decreased following inoculation from 10^8^ cells/mL to approximately 10^6^ cells/mL (Figure 5D; Supplementary figures 3A, 4A), with a corresponding decrease in MMP activity (Fig. 5F; Supplementary figure 3B, 4B). In challenged, symptomatic fragments a subsequent rise of up to 10 fold in MMP activity was regularly observed starting at 4-6 h post inoculation (Fig. 5F; Figure 6; Supplementary figure 4B). This increase was followed by an increase in bacterial load of up to half an order of magnitude at 7-10 hours post inoculation (Figure 5D; Supplementary figure 4A). These late increase in bacterial abundance and MMP activity were not observed in either the challenged, non-symptomatic fragments or in fragments challenged with *V. fischeri.* Comparing total and DsRed-labeled bacterial numbers in the outflow of one experiment revealed that the portion of DsRed labelled bacterial cells went down from 100% labeling in the inoculum to 70-90% in the following 8h, and further decrease to 60% during the subsequent rise in bacterial abundance starting at 9.5 h from inoculation (Supplementary Figure 5).

**Figure 6:**
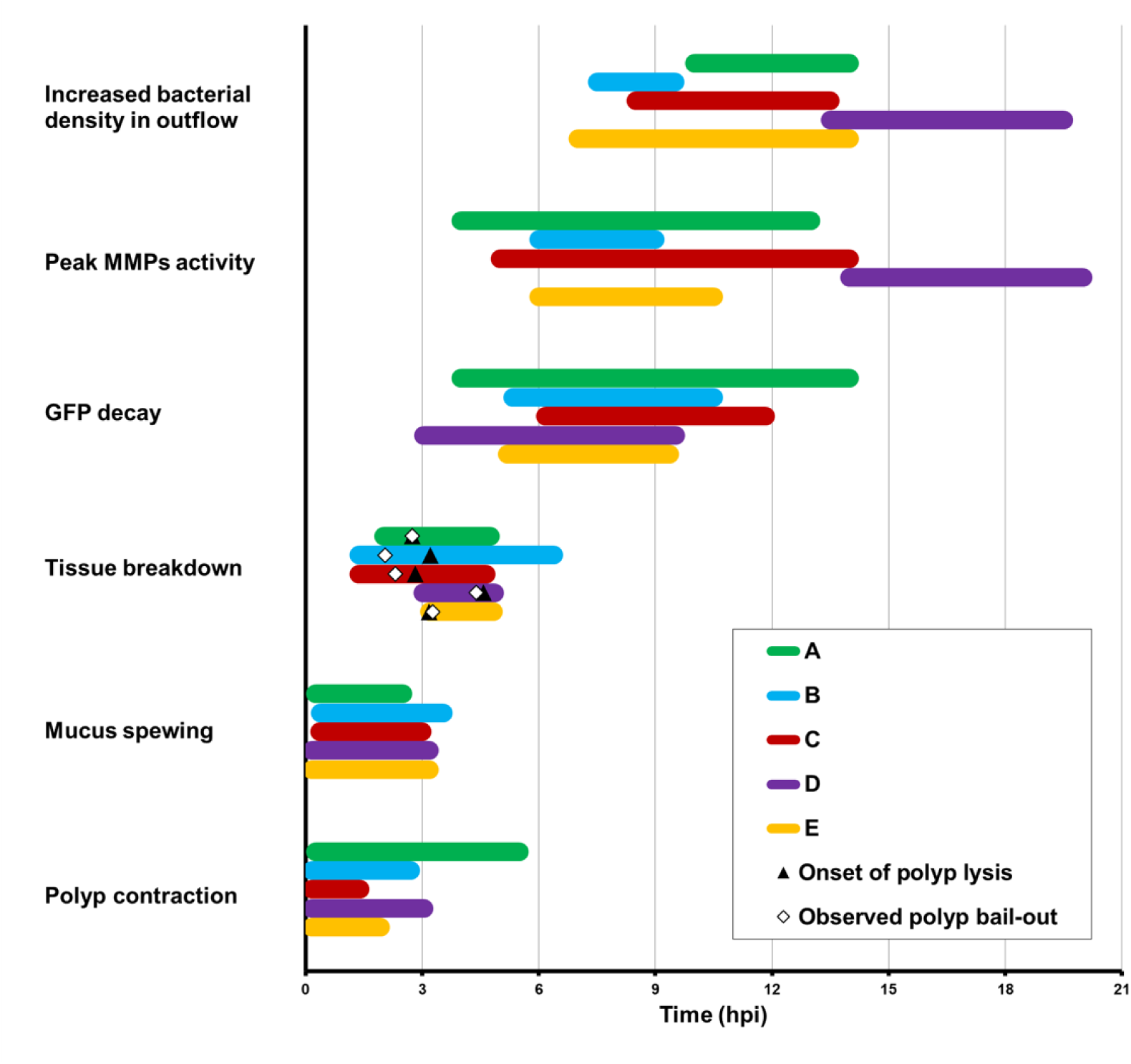
Summary of key infection phases. Results from 5 representative experiments using fragments from three *P. damicornis* colonies (A-E; see details in Supplementary Table 1). Only fragments showing clear symptoms following challenge by *V. coralliilyticus* are represented. The overall sequence of morphological, biochemical and microbial events is highly conserved although exact timing of events varied. Colored bars represent approximate duration of observation or measurement for each experiment. Discrete observations related to colony breakdown, including polyp bail-out and tissue lysis, are marked on the corresponding bar as (◊) and (▲), respectively.

By integrating the various measurement and observations from all MCI experiments into a single timeline we were able to generate the most detailed description to-date of the infection of a reef building coral by a bacterial pathogen (Fig. 6). This unified timeline reveals the conservation of the different stages described here, as well as some variability in the relative timing of specific events, particularly in the later stages. From this integrated timeline it is evident that initial polyp contraction and mucus spewing was an immediate response shared by all fragments regardless of their ultimate fate. The rise in MMP activity was always preceded by the onset of tissue lysis, and was generally preceded by GFP decay. The rise in bacterial abundance in the effluent was generally observed following the rise in MMP activity. In the majority of actively infected fragments, the entire infection process was complete within 10-15 hours following inoculation.

## Discussion

Coral disease progression is often studied at scales ranging from single colonies to the entire reef, thus overlooking the microscale processes governing host-pathogen interactions. While important insights into disease etiology have been gained by careful pathological and histological studies ^23, 27-30^, such studies are mostly limited to snapshots of the disease process at acute and morphologically recognizable phases^8, 13, 31-33^. Thus, an *in vivo* description of the sequence of microscopic events underlying the process of coral infection is still missing. The current work aims to bridge this gap, by studying coral disease under controlled laboratory conditions, at temporal and spatial scales relevant to the microscopic interactions between a bacterial pathogen and its coral host.

Using live-imaging microscopy we are able, for the first time, to visualize *V. coralliilyticus* as it colonizes and infects its coral host. Rather than colonizing the entire colony surface, we show that bacterial accumulation occurs primarily at the polyp pharynx, which points towards a gastrovascular route of infection. This is in agreement with observations reported in our previous work, demonstrating accumulation of *V. coralliilyticus* in the gastrovascular cavity of micropropagated *P. damicornis* polyps^24^. Following inoculation the majority of coral fragments displayed a clear pathology leading to colony disintegration followed by the release of *V. coralliilyticus* cells to the surrounding water. Accumulation at the coral pharynx was also observed in coral fragments challenged with *V. fischeri*, but to a lesser extent and no apparent pathology. Colonization of the colony surface in our experiments appeared to be limited to sites of tissue lesions, which may serve as hotspots of bacterial infection.

Beyond tracking of the bacterial pathogen, the MCI provides us with the unique ability to observe microscale patterns of coral behavior, a subject that is rarely considered in the context of coral disease. We were thus able to describe and characterize the sequence of behavioral responses of the coral host following a bacterial challenge. We observed specific coral reactions immediately following inoculation with pathogenic *V. coralliilyticus*, particularly the retraction of coral polyps into their calices followed by mucus spewing, that were distinct from those observed following a challenge by the non-pathogenic *V. fischeri.* Polyp retraction is a universal response of corals to physical or environmental stress^34^, indicating that the coral is sensing, and responding to, the presence of pathogens or their exudates. Moreover, as polyp retraction minimizes intake of water into the polyp gastrovascular system, and thus the internalization of planktonic food^35-37^, this behavior may also provide a means for the coral to avoid further accumulation of pathogens in its gastric cavity. The subsequent spewing of viscous, bacterial-laden mucus from the polyp mouth may be interpreted as further attempt by the coral to rid itself of the ingested pathogens, not unlike the coughing of phlegm during a throat or lung infection. Again, mucus spewing is a common coral response to high loads of food or particulate matter in surrounding water^38-40^. However, considerably less mucus was secreted in response to challenge with *V. fischeri*, suggesting that coral challenged by *V. coralliilyticus* experience stress that is not directly related to the presence of bacterial cells in the surrounding water. Thus, polyp contraction and mucus spewing may be considered an important coral defense behavior when faced with high loads of bacterial pathogens in their environment.

In all *V. coralliilyticus*-*challenged* fragments, mucus spewing was invariably followed by stretching of ceonosarc tissues. In symptomatic fragments, this ultimately led to tearing of the tissue and separation of adjacent polyps. Surprisingly, many of these isolated polyps survived, some remaining attached to the skeleton while most undergoing polyp bail-out^24, 26^. Notably, bailed out polyps collected and maintained in filtered sea-water following infection experiments remained viable for over two weeks, suggesting that these polyps were indeed able to overcome the invading pathogen. Polyp separation thus provides the coral with an additional defense layer, enabling it to quarantine disease by “sacrificing” infected polyps. This response likely prevents pathogens from spreading to the rest of the colony through the common gastrovascular system, similar to plant hypersensitive resposne^41^. Polyp bail-out may further promote the survival of the genotype by salvaging individual polyps from doomed colonies, which may settle and regenerate into new colonies where conditions are more favorable^24, 26, 42^.

A major question arising from our results relates to the function of matrix metaloproteases (MMP) in the infection sequence. MMPs were previously suggested to be a key virulence factor of *V. coralliilyticus^18^.* In our experiments, despite the exposure of all treated corals to unnaturally high levels of MMPs secreted by the bacterial pathogens during inoculation step (Figure 5B), over 30% of the corals ultimately survived the infection. Thus, under the conditions tested, MMP activity alone was not sufficient to kill the corals. This is in agreement with previous work reporting similar infectivity of a different strain of *V. coralliilyticus* following deletion of a gene encoding MMP production^12^. In our experiments we observe a rise in MMP activity at a relatively late stage of the infection, when polyps are likely already dead or dying as indicated by the decay in GFP signal. The secretion of metaloenzymes at this stage suggests their involvement in the breakdown of coral tissue, as a means for *V. coralliilyticus* to scavenge nutrients and essential metabolites from the dying colony.

An interesting observation arising from our experiments was that a large fraction of the microorganisms released in the system’s exudates over the course of the infection were not DsRed-labeled (Supplementary figure 5). While this may be explained by the loss of DsRed-encoding plasmids from transformed *V. coralliilyticus*, an alternative explanation is that additional bacterial populations, formerly part of the coral holobiont, benefit from the lysis of the coral tissue and the associated abundance of nutrients. Future analysis of the bacterial community released from corals infected under similar settings may provide further insights into the identity and nature of such rogue members of the coral microbiome.

One of our goals in constructing the MCI system was to elucidate the route of infection and disease initiation. Previous studies demonstrated involvement of motility in pathogenic *Vibrio*-coral interactions, and suggested that chemotaxis towards coral mucus facilitates host-localization and colonization of the coral surface^43-46^. This view is challenged by a recent work demonstrating increased infectivity of *V. coraliilyticus* cells with impaired chemotaxis^47^, similar to results in *V. cholera^48^* but differing from the fish pathgoen *V. anguillarum^49^.* Indeed, bacterial chemotaxis occurs over relatively short distances (100’s of microns) and requires a stable and continuous gradient of the chemoattractant^50^. As recently demonstrated, such conditions are not typically found near the surface of scleractinain corals. Ciliary flows exceeding 1 mm/s at the coral surface actively mix the coral’s bounday layer by creating vortices extending up to 2 mm into the surounding water^51^. These rapid currents, ten time the swimming speed of *V. coralliilyticus*^17^, disrupt diffusion gradients that would otherwise develop in the coral’s boundary layer, while sweeping away any pathogens reaching the coral’s surface. Thus, ciliary flows are likely to prevent pathogens of scleractinian corals from chemotaxing towards their potential hosts.

This putative role of cilia as a physical barier to bacterial colonization is further supported by the observed accumulation of pathogens at tissue lesions, where tissue confluence is breached and ciliary motion is likely disrupted. Such local patches of reduced ciliary flow, possibly enriched with infochemicals exuded from the torn tissue, may indeed facilitate bacterial chemotaxis. This may account for the rapid colonization and infection at lesion sites in our experiments. Indeed, previous studies showed that wounds caused by trauma to coral colonies provide “hot spots” for initiation of various coral diseases, including white plague, brown and black band, and others^52-54^. Here we show that even minor lesions, under the right conditions, may serve as a possible point of entry for bacterial pathogens.

The question of chemotaxis is also relevant to the accumulation of pathogens at the coral pharynx. Significantly, while pathogen accumulation at lesion sites was only observed 45-60 minutes from inoculation, comparable accumulation at the coral pharynx is observed already 10-15 minutes into the experiment, suggesting different mechanism may be driving the two phenomena. We suggest that accumulation at the pharynx may be driven by the active uptake of water into the coral’s gastrovascular system prior to polyp contraction, as part of ongoing feeding and gas exchange processes^37, 55-57^. Once inside the gastrovascular channels, where flow is likely to be laminar and boundaries within easy reach, chemotaxis may well play a part in bacterial colonization of the gastrovascular mucus.

The use of small coral fragments in our system allowed us to perform a relatively large number of experiments, using fragments from multiple colonies. An unexpected result was the heterogeneity in the response of different *P. damicornis* colonies to *V. coralliilyticus* infection (Supplementary Table 1). While some colonies were highly susceptible to infection (e.g. colonies 2 and 5), other colonies had remarkably high survival rates (e.g. colonies 3 and 4). The mechanisms underlying these differences are not clear. Genetic differences, life histories or microbiome composition may all contribute to coral resilience^58-62^. Heterogeneity was also observed at the response of individual polyps, and that too requires further investigation. Future experiments examining the genetic and epigenetic (including microbiome composition) background of different colonies may help resolve some of these questions.

It is important to note that in all bacterial challenge experiments reported here, we used approximately 10^8^ *V. coralliilyticus* cells/ml, a number that is clearly unrealistic ecologically. Inoculations with 10^7^ cells/ml or less did not result in coral mortality under the conditions tested, even following 72 hours of subsequent incubation (data not shown). While this may signify a limitation of the short duration of our experiments, it is notable that previous experiments reported for the same coral-pathogen system also used *V. coralliilyticus* concentrations of between 10^7^ and 10^8^ cells/ml to induce infection^9, 14, 46^. Ushijima and colleagues^47^ determined the infectious dose of *V. coralliilyticus* towards *Montipora* to be between 10^7^ and 10^8^ cells/ml. This suggests a compatible coral immunity and high resilience to the low densities of planktonic *V. coralliilyticus* prevalent in the reef environment^63^. An alternative route for coral infection under natural conditions, which remains to be explored, is the ingestion of vibrio-laden marine snow or infected zooplankton, delivering an infective dose of bacterial pathogens directly into the corals’ gastrovascular system^64^.

The MCI experimental setup, presented here for the first time, combines advanced live imaging microscopy, microfluidics and time resolved sampling and analysis of the system effluents enabling the evaluation of coral-pathogen interactions at unprecedented detail. This revealed several hitherto unknown aspects of coral disease, including localization of pathogens at the onset of infection, behavioral defensive responses of the coral host, and the heterogeneity of polyp fate following infection, and defined distinct phases of the infection process. Future application of approaches similar to that described here will facilitate more detailed understanding of the complex and ecologically important interactions occurring between corals and their bacterial pathogens. This platform will provide a foundation for future studies aiming at elucidating the versatile adaptive response of the fragile coral ecosystems to fluctuating environments.

## Methods

### MCI experimental setup

Microfluidic chambers were fabricated *in*-*house* as follows: A 5×1.5 cm slab was cut out of a 5 mm thick sheet of polydimethylsiloxane (PDMS) silicone elastomer (Sylgard^®^ 184) using a utility knife. 4-6 Ø8mm wells were punched into the resulting slab using a biopsy punch of the same diameter, forming chambers of approximately 250μL. Inlet and outlet holes were punched into opposing sides of each chamber using a 1 mm biopsy punch (Integra^®^, Fischer Scientific). (Supplementary figure 1). The PDMS slab was then bonded to a glass microscope slide by exposing both to oxygen plasma for one minute using a laboratory Corona Treater (Electro-Technic Products). Each chamber was fitted with polyethylene inlet and outlet tubing (BPE-60, Instech Laboratories) (Fig 1). The assembled device was placed on a temperature controlled microscope stage. Small *P. damicornis* fragments (3-5 mm) were placed in each chamber and chambers sealed with ApopTag^®^ Plastic cover slips (Merck). Flow (2.6 mL hour^-1^) was generated using a peristaltic pump (Ismatec) connected to the outlet tube. The input tube was connected to a flask containing FASW (0.22 μm). Inoculation was carried out by transferring the free end of the inlet tube to a flask containing FASW supplemented with 10^8^ of either *V. coralliilyticus* or *V. fischeri* for 2 h, and then transferring back to the FASW-containing flask.

The outlet stream from each chamber was continuously collected using a 4 channel fraction collector (Gilson Inc.) into 2 ml Eppendorf tubes, with tubes for each stream changed at 30 min intervals. Tubes were maintained in an aluminum tube rack placed in an ice bath to maintain contents at 0-1°C. Every 2^nd^ tube was supplemented in advance with paraformaldehyde (PFA) to a final concentration of approximately 1%, enabling subsequent bacterial quantification using flow cytometry. Fractions from tubes without fixative were centrifuged, and supernatant used for quantification of MMP enzymatic activity.

### Coral collection and handling

All *P. damicornis* colonies used in this study were collected from a coral nursery located at a depth of 8 m off the pier of the Inter-University Institute, Eilat, Israel (Israel nature and parks authority permit No # 2014/40327). Collected corals were maintained in an aquarium at the Weizmann Institute of Science. Small branch tips were clipped from the colonies and left in the main tank for recovery for at least one week. Prior to each experiment some fragments were transferred to a separate 4 L tank filled with FASW and incubated at 31°C for a period of 3 days. The fragments were then transferred to the MCI device for microscopic observation. At the beginning of each experiment, prior to inoculation, fragments were acclimated on the stage for at least 3 hours with a constant flow of filtered aquarium water.

### *Vibrio coralliilyticus* transformation to express DsRed

Infection experiments were performed using the *V. coralliilyticus* strain YB2 labelled with a plasmid encoding for a potent variant of DsRed2 fluorescent protein^25^ as described previously^24^. For each experiment, DsRed-labelled *V. coralliilyticus* were grown overnight from glycerol stock at 30 °C in Zobell Marine Broth. Bacteria were then centrifuged (3500 G, 5 minutes) and resuspended in FASW. Tubes were then incubated at 30°C with no shaking to allow sinking of non-motile bacteria.

### Infection assays procedure

#### Experimental procedure

The general work flow and infection scheme is illustrated in **Error! Reference source not found.**. Inoculation was carried out by flowing a suspension of DsRed-labeled *V. coralliilyticus* or *V, fischeri* (approximately 10^8^ cells/mL) into the chamber over a period of two hours. Inlet flow was then switched to filtered aquarium water for the remaining incubation. Live imaging microscopy was carried out using a fully motorized inverted fluorescence microscope (Olympus IX81) equipped with a Coolsnap HQ2 CCD camera (Photometrics). Throughout the infection experiments, multichannel micrographs of the fragments were captured every 15 minutes at 4X magnification. This enabled visualization of the coral-tissue GFP, zooxanthellae chlorophyll, and DsRed fluorescence, alongside a bright-field channel.

### Image analysis

Image analysis was carried out using imageJ (FIJI), by measuring mean grey intensity (in pixels) of the entire frame in each channel (GFP, Chlorophyl, and DsRed) captured at every time point.

### Downstream exudate analysis

To couple the visual observations with direct microbial and biochemical measurements, each chamber’s effluents was continuously collected in a time resolved manner using a fraction collector and immediately cooled to between 0 and 2°C (Figure 1, step 5). Odd numbered fractions (1.3 mL) each were immediately fixed in 1% PFA in FASW. 20 μL of each sample was diluted 10 fold and stained with nucleic acid stain SYBR-gold (Invitrogen). Cell abundance was measured using a flow cytometer (iCyt Eclipse, excitation: 488 nm, emission: 500–550 nm). Even numbered fractions were collected with no fixation and filtered through 0.22 μm syringe filter (Millipore). Filtrate was used to estimate MMP activity using a specific fluorescent substrate (Calbiochem MMP-2/MMP-7 Substrate, Fluorogenic) in a microplate reader (Tecan Infinite^®^ M200pro). Fluorescence (exitation :325 nm, emmition: 393 nm) was measure every 90 seconds at 30°C for 40 min. We used a specific MMPs substrate to demonstrate that isolated *V. coralliilyticus* secretes MMPs to the culture medium during different growth phases (Supplementary figure 6A). This specific activity was confirmed by inhibition with GM6001, a broad-spectrum MMPs inhibitor with inhibition capacity of IC_50_ = 5 μM (Supplementary figure 6B).

## Footnotes

**Author contributions** ARG and OHS developed the MCI experimental setup. ARG transformed *V. coralliilyticus* and performed the experiments and image analysis. EKW contributed to experimental design and interpretation of results. AV contributed to the design and evaluation of the experiments and the overview of all aspects of the project. ARG, OHS, EKW, and AV wrote the manuscript.

## Acknowledgements

*V. coralliilyticus* transformation was carried out with the help of D. Schatz using materials and protocols generously provided by Prof. E. Stabb (Franklin College, University of Georgia). Inna Solomonov and Irit from the Irit Sagee lab, Weizmann Institute of Science, are assisted with establishment and validation of the MMP activity essay. This work was supported by the Human Frontiers in Science Program (award #RGY0089), the Weizmann - EPFL Collaboration Program (grant number: 721236), and the Angel Faivovich Foundation for Ecological research.

